# A step towards animal models with self-sustained fungal bioluminescence

**DOI:** 10.1101/2025.08.04.668132

**Authors:** Hannah J Gleneadie, Nicolas Veland, Eleanor C Warren, Alessandro Sardini, Consuelo Barroso, Anastasia A. Fadeeva, Zoe Webster, Dirk Dormann, Feng Gao, Enrique Martinez-Perez, Matthias Merkenschlager, Karen Sarkisyan, Amanda G Fisher

**Author notes:** These authors contributed equally (Karen Sarkisyan, Amanda G Fisher).

## Abstract

Engineering animal models with self-sustained luminescence could enable non-invasive longitudinal monitoring of molecular events in living animals. To create animal models that report physiology with autoluminescence, both luciferin biosynthesis enzymes and the luciferase need to be optimised. Previous work on engineering the autoluminescence pathway from fungi resulted in the development of nnLuz_v3, a version of *Neonothopanus nambi* luciferase with enhanced thermal stability. Here, we generated an nnLuz_v3 reporter of endogenous *Cyp1a1* expression as a measure of aryl hydrocarbon receptor (AHR) activation, assessing the performance of nnLuz_v3 *in vivo* at physiologically relevant expression levels. As AHR dynamically responds to metabolic, environmental and dietary changes it provides a validated platform to assess novel luminescence approaches. In *Cyp1a1-nnLuz* mice bioluminescence signal was stable, allowing the generation of well-resolved luminescence images both on standard *in vivo* imaging equipment and consumer-grade cameras. Using mice and nematode models, we demonstrated limited oral availability of the fungal luciferin, potentially compatible with delivering the substrate via food or the microbiome. Our results are an encouraging first step in the generation of an autoluminescent mammalian model of a molecular event and encourage optimisation of other enzymes of the fungal luciferase pathway.

---

Light is the most commonly used modality for reporting molecular events in living organisms. Yet, its use for deep tissue imaging in animals is limited by scattering and absorption by pigments which interfere with light propagation. Under these conditions, fluorescence-based molecular reporters become largely unusable due to autofluorescence of tissues and the poor delivery of excitation light^1^. Luminescence reporters possess the advantage of virtually no background signal, but require exogenous addition of the luciferin substrate ^2,3^. This makes imaging invasive, discontinuous, expensive and subject to uneven tissue distribution of the substrate.

For a number of *in vivo* applications, the ultimate light-based reporter technology could be self-sustained luminescence. When, in a single organism, the luminescence substrate is produced biosynthetically and the luciferase enzyme used to report on a molecular event, obtaining direct physiological information becomes as simple as taking a photo in the dark. Two pathways for luciferin biosynthesis have been fully deciphered, the bacterial (lux) pathway ^4^ and the fungal (luz) pathway ^5^. There has been significant progress in adapting these to function in plants and mammalian cells ^6-11^, leading to the recent proof of concept demonstration of autoluminescent mice which constitutively express the complete bacterial bioluminescence pathway ^12^. However, the emission of blue-green light by the luxAB luciferase is not optimal for *in vivo* imaging, perhaps contributing to the uneven distribution of luminescence and the modest light output observed ^12^. Moreover, constitutive, parallel expression of both the luciferase and the enzymes required for luciferin biosynthesis limits the usefulness of this system. In this study, we aimed to derisk expression of the fungal bioluminescence pathway in animals by first assessing whether the fungal luciferase nnLuz can be used as a reliable molecular reporter.

The fungal bioluminescence pathway is a short cycle within fungal phenylpropanoid metabolism, catalysed by four enzymes, with the light-emitting reaction performed by the luciferase, Luz (Figure 1a). Heterologous expression of these enzymes in plants and fungi allowed generation of organisms with self-sustained luminescence ^8^. Previous engineering efforts produced nnLuz_v3, a version of the luciferase with enhanced stability at elevated temperatures ^9^. We first tested nnLuz_v3 in a mammalian cell line. We expressed the luciferase in HEK293T cells under control of the strong CAG promoter as a T2A-separated fusion gene with the far-red fluorescent protein iRFP670. We observed stable luminescence and fluorescence signals and no abnormal cell phenotypes or toxicity (Supplementary Figure 1).

**Figure 1:**
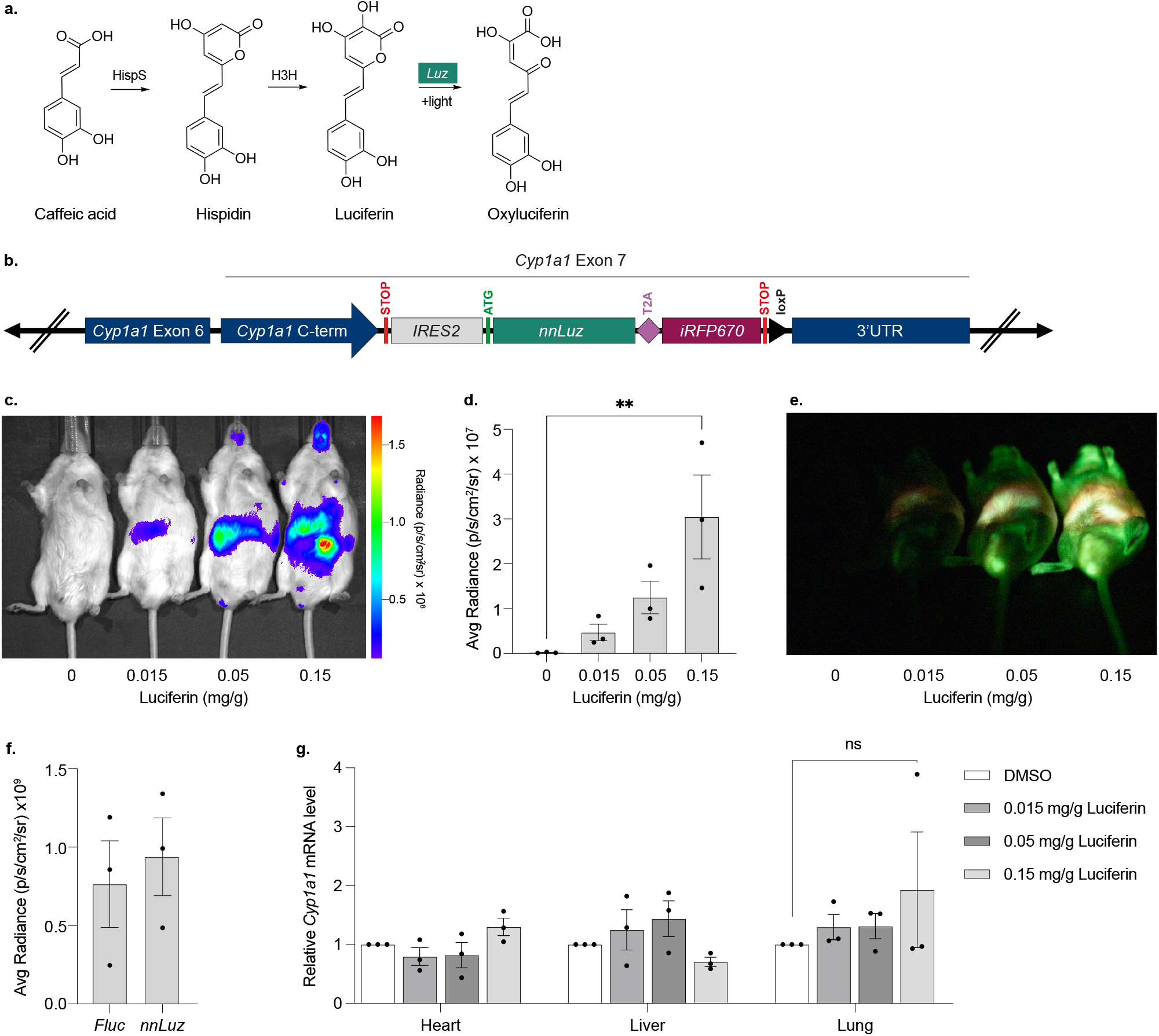
Generation and characterisation of an endogenous bioluminescent reporter of AHR activation using the fungal luciferase *nnluz_v3*. **a**. Schematic outlining the fungal luciferin pathway. Hispidin synthase (HispS) converts caffeic acid into hispidin which is then hydroxylated by H3H into 3-hydroxyhispidin (fungal luciferin). The luciferase enzyme (Luz) converts luciferin into oxyluciferin and produces light ^5^. **b**. Schematic outlining the engineered *Cyp1a1* locus in *Cyp1a1-nnLuz* mice and mouse embryonic stem cells (mESCs). Fungal luciferase (*nnLuz_v3*) and fluorescent protein, *iRFP670*, were inserted before the 3’UTR of endogenous *Cyp1a1*. An *IRES2* sequence separates the reporter genes from the *Cyp1a1* C-terminal and a *T2A* sequence separates the two reporter genes. The targeting strategy is outlined in supplementary figure 2. **c**. Representative bioluminescent image of *Cyp1a1-nnLuz* adult mice 5 h post intraperitoneal (IP) injection with 3mC. IP injection of fungal luciferin at concentrations of 0, 0.015, 0.05, and 0.15 mg/g show dose-dependent luminescence. **d**. Quantification of **c**. Graph shows mean average radiance of whole mice (n=3) +/- SEM. A one-way ANOVA compares vehicle with luciferin-injected animals with Dunnetts multiple comparison test. ^**^ p<0.01. **e**. The same animals from **c**. imaged using a Sony Alpha 1 camera with SEL50F12GM lens, ISO 800 and exposure of 300 secs. **f**. *Cyp1a1-Fluc* and *Cyp1a1-nnLuz* mice were IP injected with D-luciferin or fungal luciferin, respectively, at 0.15 mg/g and imaged across a 90-minute time-period (full details shown in supplementary figure 4). The graph shows average radiance emitted across this period. Mean (n=3) +/- SEM. All mice were IP injected with 3mC 5 h prior to imaging. **g**. Heart, liver, and lung tissues dissected from *Cyp1a1-nnLuz* mice 1 h after fungal luciferin injection. Relative *Cyp1a1* mRNA levels as a measure of AHR activation. Bars show mean (n=3) +/- SEM with a two-way ANOVA to compare vehicle to luciferin treated samples and Dunnetts test to correct for multiple comparisons.

We then aimed to assess nnLuz_v3 as a physiology reporter, using aryl hydrocarbon receptor (AHR) activation in mice as a model. AHR is a soluble transcription factor that recognises numerous endogenous and environmental ligands, adjusting cellular physiology to maintain tissue homeostasis^13^. One of its downstream effectors is the cytochrome P450 protein Cyp1a1. Activation of *Cyp1a1* expression is a reliable reporter of AHR activation ^14,15^. A mouse model that translates AHR activation into autonomous light emission would enable new types of non-invasive experiments to continuously monitor toxic exposures and autoimmune responses. As a step towards generation of such a model, we designed an IRES-driven *nnLuz_v3-T2A-iRFP670* cassette for integration into the 3’UTR of endogenous *Cyp1a1* in the mouse genome (Figure 1b, Supplementary Figure 2). Integration of the cassette into mouse embryonic stem cells resulted in multiple knock-in clones that demonstrated activation of green luminescence upon treatment with the AHR agonist FICZ (Supplementary Figure 3). Clone A4, which demonstrated a strong bioluminescent signal, whilst remaining morphologically unchanged from the parental mESC line, was used to generate transgenic mice.

Transgenic animals appeared to develop normally and showed no apparent phenotypic changes. Pre-treatment of animals with long-acting AHR agonist 3-methylcholanthrene (3-mC) led to development of robust luminescence signal, with brightness dependent on the concentration of intraperitoneally (IP) injected luciferin (Figure 1c-e). The overall signal intensity was similar to that of an equivalent *Cyp1a1* reporter line expressing firefly luciferase ^16^ (Figure 1f) but showed different kinetics, with luminescence from Fluc achieving higher peak signal intensity, at 10-15 minutes post luciferin injection, while nnLuz was more stable across the time-course examined (Supplementary Figure 4). The brightness was sufficient for imaging on consumer-grade cameras with 50 megapixel resolution (Figure 1e). Encouragingly, we observed no acute toxicity upon injection with the fungal luciferin and at concentrations required for imaging, luciferin itself did not induce AHR activation (Figure 1g).

Interestingly, while bioluminescence from nnLuz is green in fungi, with the maximum emission at around 520 nm, in mice, luminescence peaked at 520 nm, but had an additional, smaller peak at ∼620 nm (Supplementary Figure 5 a-b). *Ex vivo* analysis of luminescence from different organs showed that light emitted from most tissues was green, with the exception of lungs that exhibited the 520/620 nm double-peaked spectrum (Supplementary Figure 5 c). As expected, both *in vivo* and *ex vivo* the firefly luciferase in *Cyp1a1-Fluc* reporter animals emitted light in the red spectrum (Supplementary Figures 5 a, b, d) while we previously detected a similar double-peaked spectrum in a green click beetle luciferase (CBG99Luc) expressing-mouse line, which again could be localised to the lungs ^17^ (Supplementary Figure 5 e). One interpretation of this result is that it is not specific to the chemistry of fungal bioluminescence nor related to the expression of *Cyp1a1*, but likely as an artefact of green-emitting luciferase enzymes, potentially attributed to the re-emission of luminescence by fluorescent molecules present in animal tissues, such as porphyrins, that are present in large quantities in lungs and fluoresce at around 620 nm ^18,19^.

We demonstrate that fungal luciferin is orally bioavailable, potentially allowing minimally invasive experiments with animals expressing nnLuz (Supplementary Figure 6,7). Mice fed fungal luciferin by oral gavage (OG) show abdominal luminescence signal from as early as 8 minutes post feeding (Supplementary Figure 6 a). However, the signal detected following OG delivery of luciferin differed both in intensity and distribution from the signal detected following IP delivery (Supplementary Figure 6 b-e). Following OG delivery luminescence is primarily detected in the intestines, with only very low signal detected in the lungs, heart or liver (Supplementary Figure 6 b, c). This is likely due to retention of the luciferin in the digestive tract following OG delivery and again highlights the potential benefits of generating a self-sustaining luminescent model.

To investigate whether fungal luciferin exhibits oral availability in an alternative animal model, we generated a transgenic *C.elegans* line expressing nnLuz under the control of the ubiquitous *Peft-3* promoter (Supplementary Figure 7 a-b). Luminescent nematodes can be used for high throughput experiments such as genetic or compound screening. Upon exposure to luciferin, the worms displayed observable bioluminescence. Microscopy analysis of bioluminescence patterns showed that luminescence was primarily localised to the pharynx, indicating that luciferin does not permeate through the cuticle and instead reflects limited oral availability (Supplementary Figure 7 c). However, as no signal was observed from the rest of the worm, it is likely that the pharyngeal-intestinal valve serves as a barrier to systemic uptake. Interestingly, nnLuz luminescence is green in nematodes with emission peaking around 530 nm, suggesting the dual peaked emission spectrum seen in mice does not translate across species (Supplementary Figure 7 d).

The generation of reporter animal models which luminescence without the addition of exogenous substrate would be an exciting new tool for the research community: current bioluminescent models (reviewed in ^20,21^) would become less invasive and the potential to split the pathway across cells or species could be harnessed to investigate host-microbe interactions (Figure 2). The fungal bioluminescence system is a promising biochemical pathway to engineer autonomous bioluminescence in animals, however, the wild type fungal luciferase, nnLuz, showed poor activity at temperatures above 30°C ^5^. Our previous study reported nnLuz_v3, a variant carrying three substitutions, T99P, T192S and A199P, that showed higher activity at the elevated temperatures required for imaging in warm-blooded animals ^9^. It was not clear whether this variant would provide a high enough signal-to-noise ratio in transgenic animals. Here, we demonstrate that nnLuz_v3 can be used as an *in vivo* reporter in mice and nematodes, and can be used to visualise inducible transcriptional activation, as shown with the AHR pathway. Longitudinal imaging of AHR activity in mice provides us with an unrivalled platform to test the applicability of luminescence approaches to model dynamic, reversible, systemic and organotypic responses to the lived environment. Encouragingly, *Cyp1a1-nnLuz* mice display bioluminescence intensity and signal distribution comparable to that of previously validated *Cyp1a1-Fluc* animals ^16^. Therefore, this work provides a promising first step in the pathway towards generating animal models with self-sustained fungal bioluminescence to visualise molecular events in real-time.

**Figure 2:**
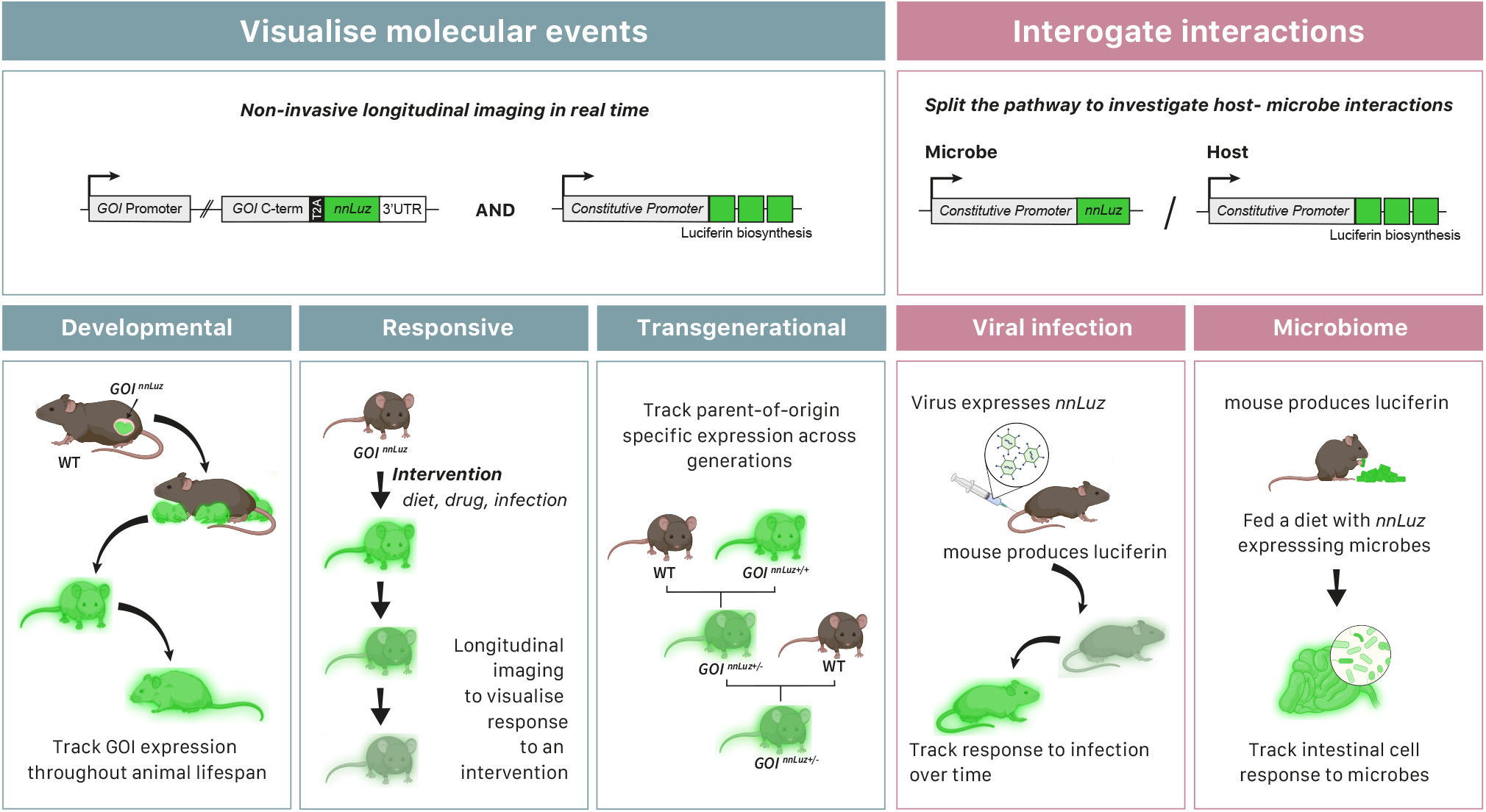
Potential applications of autoluminescent mice. Schematic representing some of the potential uses of animal models expressing the fungal luciferase/luciferin pathway. **Visualise molecular events:** The first panels (blue) show the use of the fungal pathway to generate molecular markers. The luciferase can be inserted into an endogenous gene of interest (GOI), while the luciferin is constitutively and ubiquitously expressed in the same animal. Therefore, the animal will luminescence whenever the GOI is expressed. This can be harnessed to improve upon developmental, responsive and transgenerational studies. **Developmental**: Non-invasive imaging allows for GOI expression to be tracked in the same animal across the entire lifespan, from *in utero* ^22^ to old age. **Responsive**: Expression of a GOI in response to an external intervention such as drug treatment, diet, environment, illness or infection. Animals can be imaged repeatedly following intervention. **Transgenerational**: Parent-of-origin specific expression can be tracked across generations, and in response to interventions, to investigate imprinting, X-linked or mitochondrial disorders. **Investigate interactions:** Second panels (pink) show the potential to split the pathway across organisms. Here a host animal would express the genes required for luciferin biosynthesis and a microbial species would express the luciferase, *nnLuz*. **Viral infection**: The virus expresses the luciferase and the mouse produces the luciferin. Non-invasive imaging could allow you to track progress of the infection in real time. **Microbiome:** Introduce *nnLuz* expressing microbes into the microbiome of a luciferin producing mouse. The tissue specific response to the microbes could be tracked.

## Acknowledgements

This work was funded by the Medical Research Council (MRC) (A.G.F. by MC_PC_23024 and MC_UP_1605/12, K.S by UKRI MC-A658-5QEA0). KSS is supported by UKRI Biotechnology and Biological Sciences Research Council through the International Science Partnerships Fund (ISPF), grant number UKRI249. AAF is supported by RSF, project number 24-74-10105 (https://rscf.ru/project/24-74-10105). N.V. received an ERDA award from the Institute of Clinical Sciences, Imperial College London.

## Author contributions

N.V., K.S and A.G.F conceptualised the study. The majority of experiments were performed by H.J.G and N.V with help from A.S, E.C.W, F.G. *C.elegans* experiments were performed by E.C.W, C.B.G and E.M.P. AAF synthesised fungal luciferin for experiments in this project. Mouse and *C.elegans* imaging support was provided by A.S and D.D respectively and mouse transgenics and husbandry support was provided by Z.W and M.M. K.S and H.J.G wrote the manuscript, with input from all co-authors.

## Competing interests

Karen Sarkisyan is a shareholder of Light Bio, Inc. All other authors declare no competing interests.

## Data availability statement

All numerical source data is provided in Supplementary Data file 1.

**Supplementary Figure 1:**
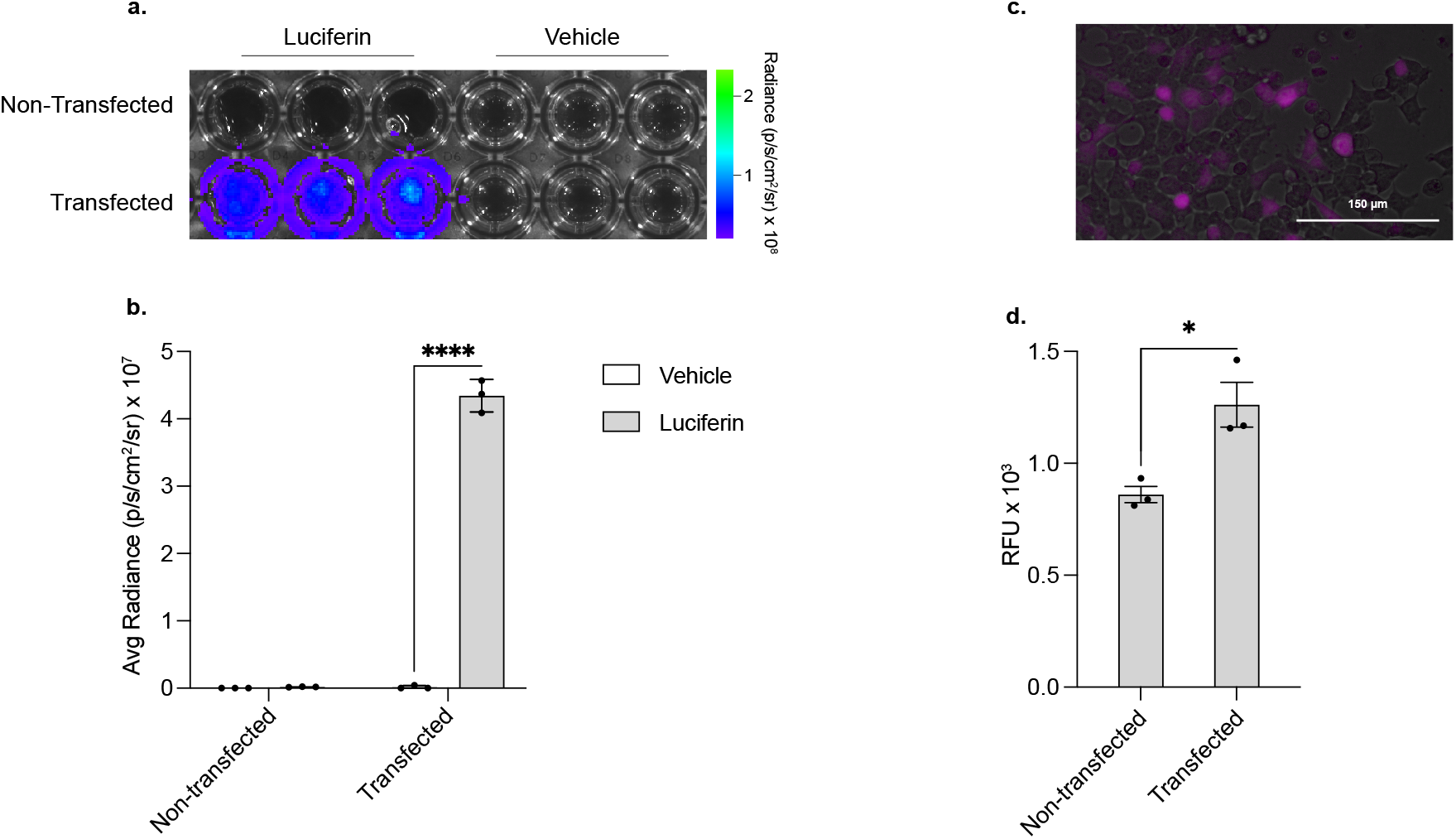
Expression of fungal luciferase in mammalian cells. Fungal luciferase *nnLuz_v3* was expressed in mammalian HEK 293T cells under control of CAG promoter through cloning as a bicistronic expression vector together with *iRFP670* separated by T2A peptide (pCAG-nnLuz-T2A-iRFP670). Detection of bioluminescence and fluorescence was performed 24 hours after transfection. **a**. Bioluminescence detection after incubation with fungal luciferin (650 µg/mL) of transfection compared to untreated cells. Transfections reactions were performed in triplicates. **b**. Quantification of fungal luciferase nnLuz_v3 bioluminescent signal as average radiance in each well from **a**. Two-way ANOVA with Šidák’s multiple comparison test to compare vehicle to luciferin treated samples, ^****^ p<0.0001. **c**. Fluorescent microscopy detection of iRFP670 in HEK 293T cells expressing pCAG-nnLuz-T2A-iRFP670. **d**. Quantification of data from **c**. Paired t-test to compare transfected with non-transfected cells, ^*^ p<0.05.

**Supplementary Figure 2:**
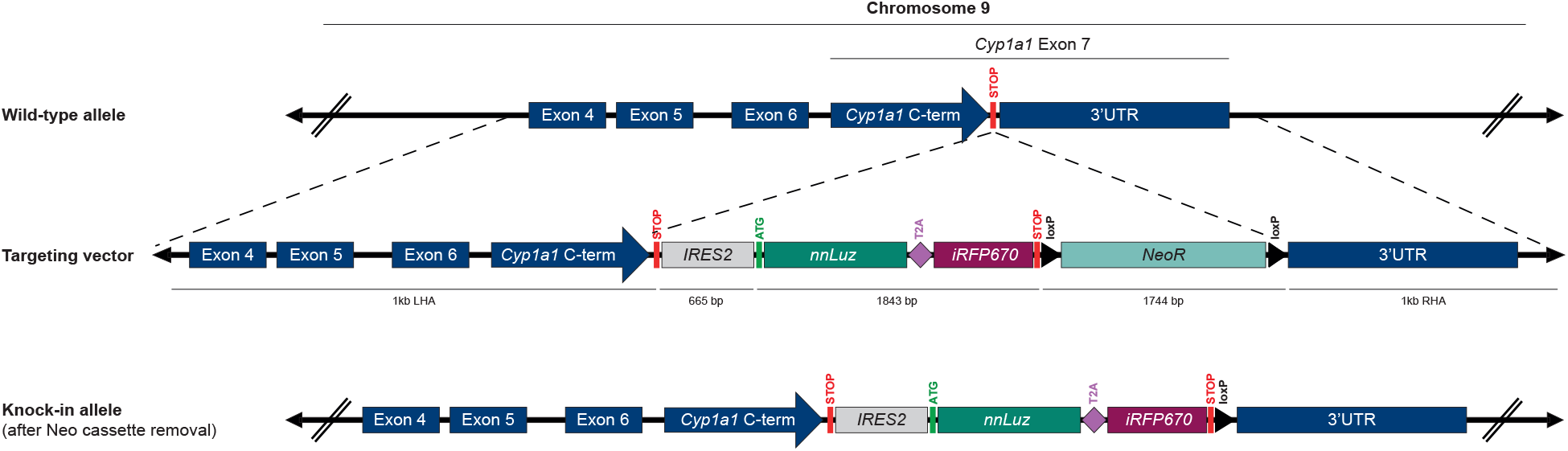
Generation of *Cyp1a1-nnLuz* reporter mESCs and mouse line. To generate the knock-in *nnLuz_v3-T2A-iRFP670* reporter into the endogenous mouse *Cyp1a1* gene, we performed gene targeting using CRISPR in mESCs with a sgRNA sequence specific to cleave DNA in the vicinity of the Stop codon in exon 7. The targeting vector used as donor DNA template contained 1 Kb of each homology arm. The left homology arm (LHA) spans exons 4 to 7 of *Cyp1a1* gene, while the right homology arm (RHA) extends across the 3’ UTR. We also included a modified version of the internal ribosomal entry site sequence (IRES2) between the stop codon of *Cyp1a1* and the start codon of *nnLuz_v3* to allow independent protein synthesis of the reporter nnLuz_v3-T2A-iRFP670 polypeptide from the endogenous Cyp1a1 protein. In addition, a neomycin resistance cassette (*NeoR*) flanked by loxP sites was included in the targeting vector for selection of successfully targeted mESC clones.

**Supplementary Figure 3:**
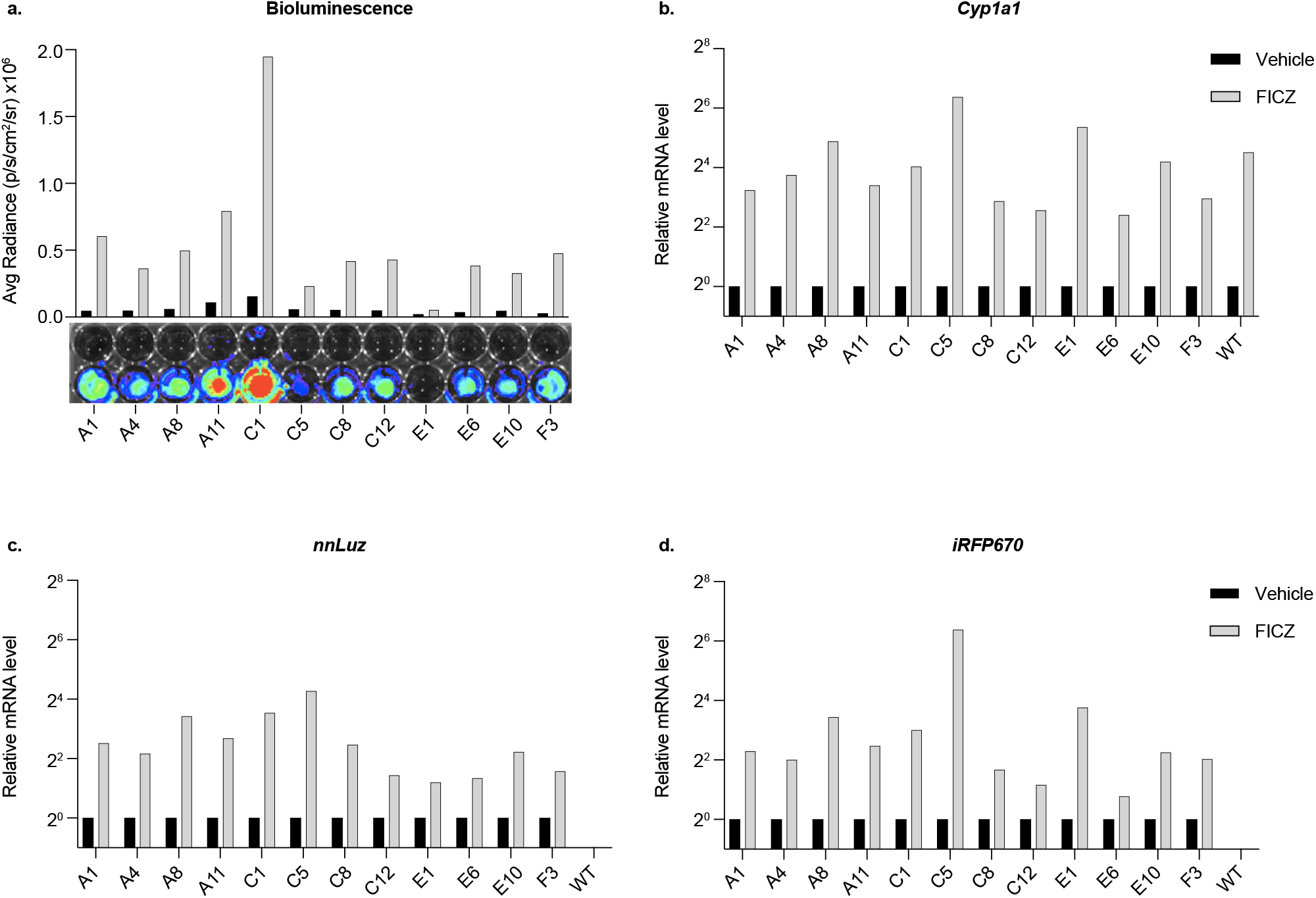
Luminescence and reporter gene expression in *Cyp1a1-nnLuz* mESCs. Multiple mESC *nnLuz_v3-T2A-iRFP670* knock-in (KI) clones were examined to determine levels of nnLuz_v3 reporter activity and gene expression in response to AHR pathway activation, to identify an optimal KI clone to generate the *Cyp1a1-nnLuz* mouse line by blastocyst microinjection. **a-d**. *Cyp1a1-nnLuz* mESC clones were exposed to FICZ or vehicle for 4 h. **a**. Quantification (average radiance) (upper), and representative images (lower), of bioluminescent signal after incubation with fungal luciferin (650 μg/mL). **b-d**. Quantification of mRNA transcripts by RT-qPCR for *Cyp1a1* (**b**), *nnLuz* (**c**) and *iRFP670* (**d**). *Gapdh* mRNA level was used for normalization of each gene target.

**Supplementary Figure 4:**
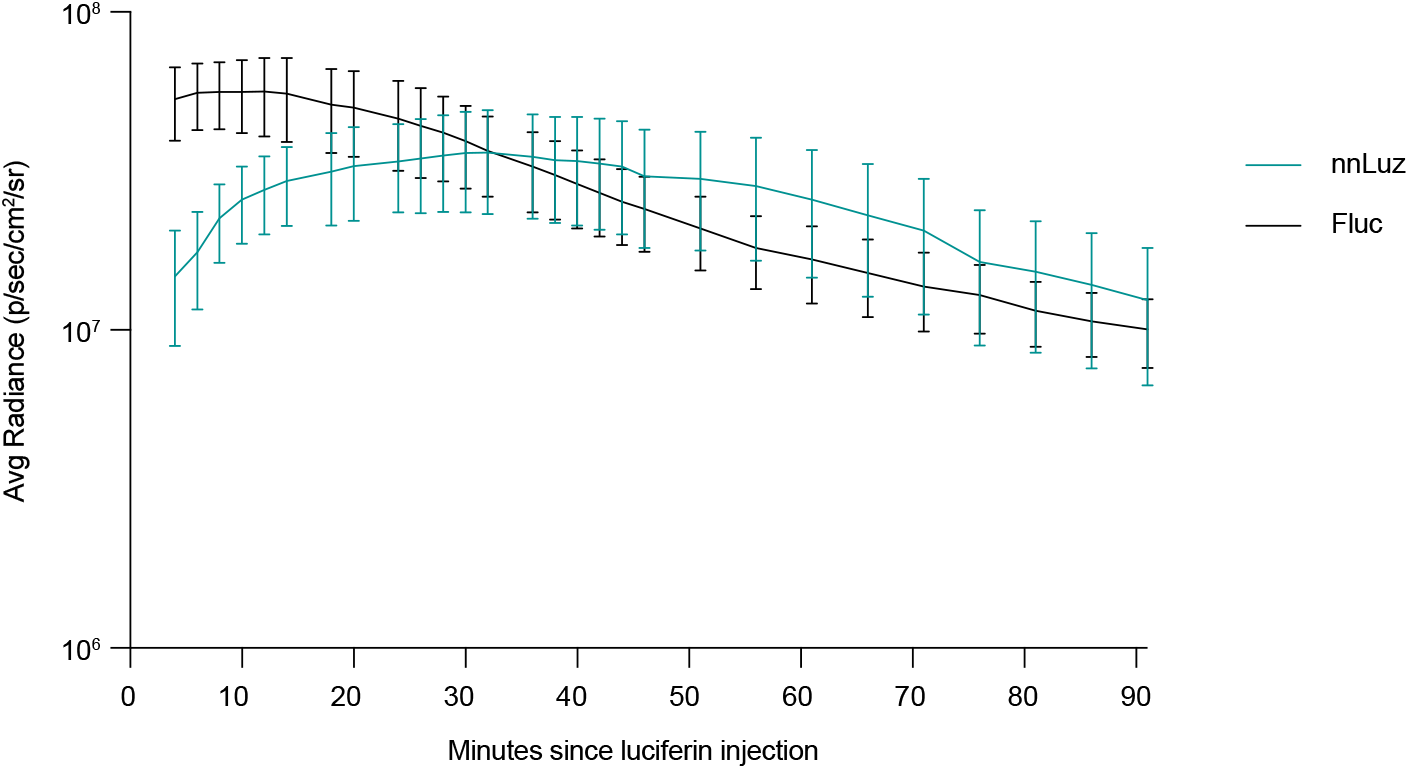
Kinetics of fungal luciferin and D-luciferin bioluminescence. Quantification of average luminescence emitted from *Cyp1a1-nnLuz* and *Cyp1a1-Fluc* mice imaged repeatedly across a 90-minute time period following IP injection with 0.15 mg/g fungal luciferin or D-luciferin, respectively. Mice were IP injected with 3mC 5 h prior to the start of imaging. Graph shows mean whole-body radiance (n=3) for each time point +/- SEM.

**Supplementary Figure 5:**
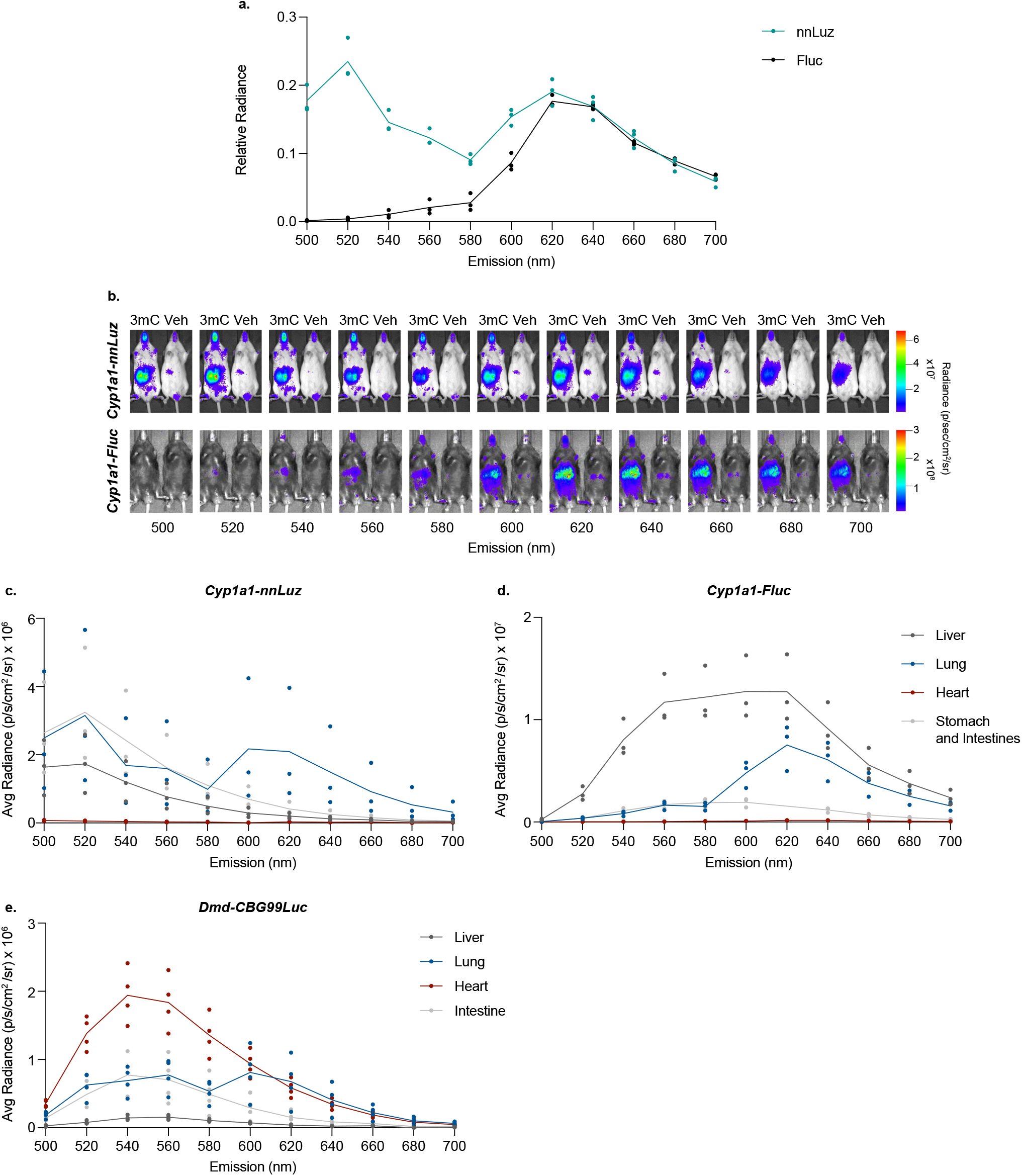
*In vivo* and *ex vivo* emission spectra of nnluz. **a-b**. *Cyp1a1-nnLuz* (nnLuz) and *Cyp1a1-Fluc* (Fluc) adult mice were IP injection with 0.15 mg/g fungal luciferin or D-luciferin, respectively, and bioluminescence imaged using 20 nm band pass emission filters across a 500–700 nm range. **a**. For each wavelength, whole-body radiance is normalised against the corresponding open-filter value. All mice were IP injected with 3mC 5 h prior to imaging. Graph shows mean (n=3). **b**. Representative images of **a**. Mice were IP injected with vehicle or 3mC 5 h prior to luciferin exposure, for each genotype and emission filter a vehicle and 3mC treated mouse are shown. **c-d**. Tissues were dissected from *Cyp1a1-nnLuz* (**c**), *Cyp1a1-Fluc* (**d**) and *Dmd-CBG99Luc* (**e**) mice after IP injection with fungal luciferin (**c**) or D-luciferin (**d-e**). Bioluminescence imaging performed using 20 nm band pass emission filters across a 500–700 nm range. Graphs show quantification of average radiance (n=3-4). **c-d**. Mice were injected with 3mC 5 h prior to luciferin exposure. For **e**. data is taken from ^17^.

**Supplementary Figure 6.**
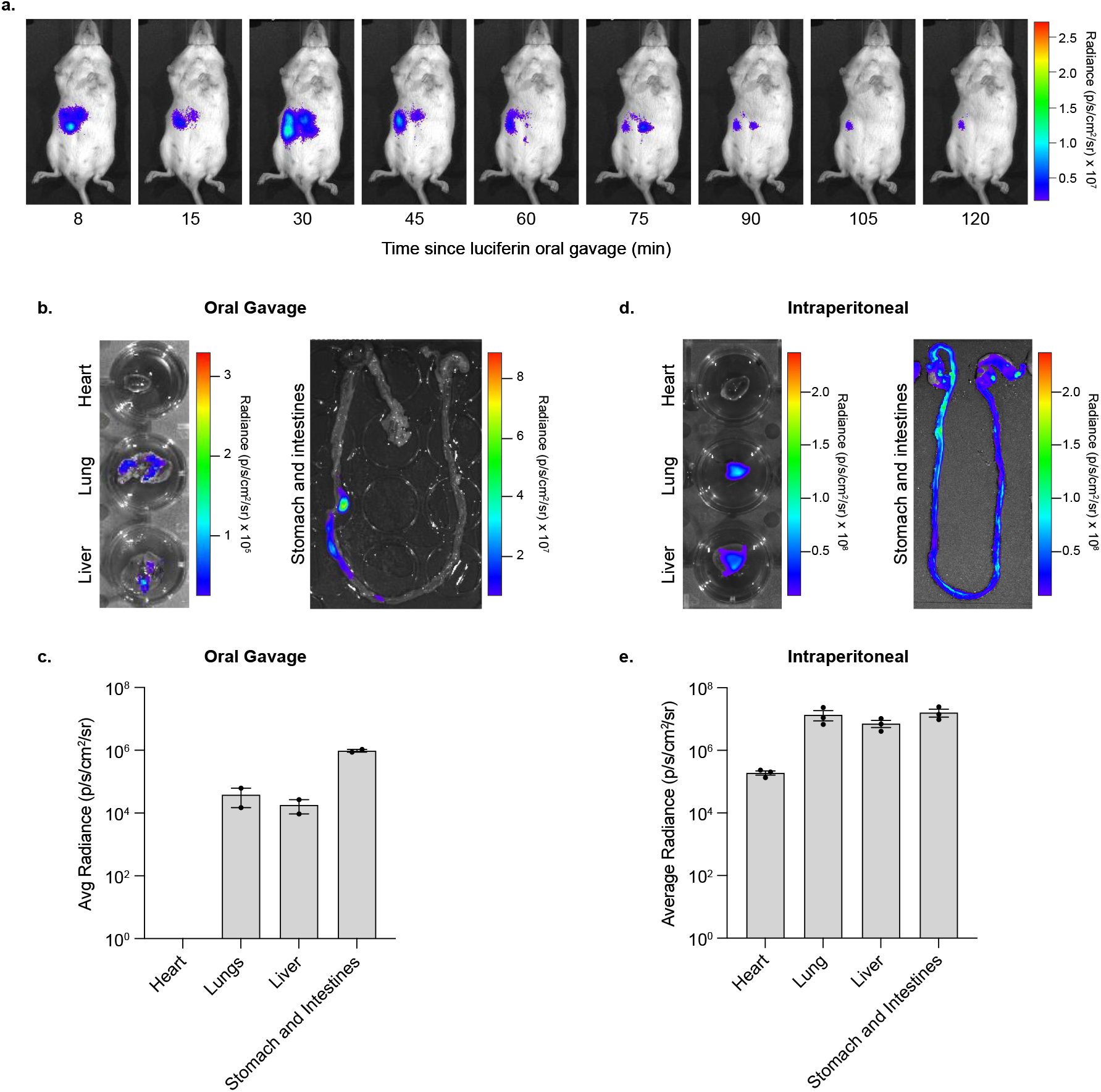
Oral availability of fungal luciferin. **a-c**. *Cyp1a1-nnLuz* mice were fed 0.15 mg/g fungal luciferin by oral gavage. Mice were IP injected with 3mC the night before imaging. **a**. Representative bioluminescence images of mice up to 120 mins post oral gavage. **b-c**. Bioluminescence imaging was performed on tissues dissected from the mice in **a. d-e**. Bioluminescence imaging was performed on tissues dissected from *Cyp1a1-nnLuz* mice following IP injection of 0.15 mg/g fungal luciferin. **b/d**. Representative images. **c/e**. Quantification of average radiance minus background radiance. Graphs are shown in log10 scale and show mean (n=2/3) +/- SEM.

**Supplementary Figure 7:**
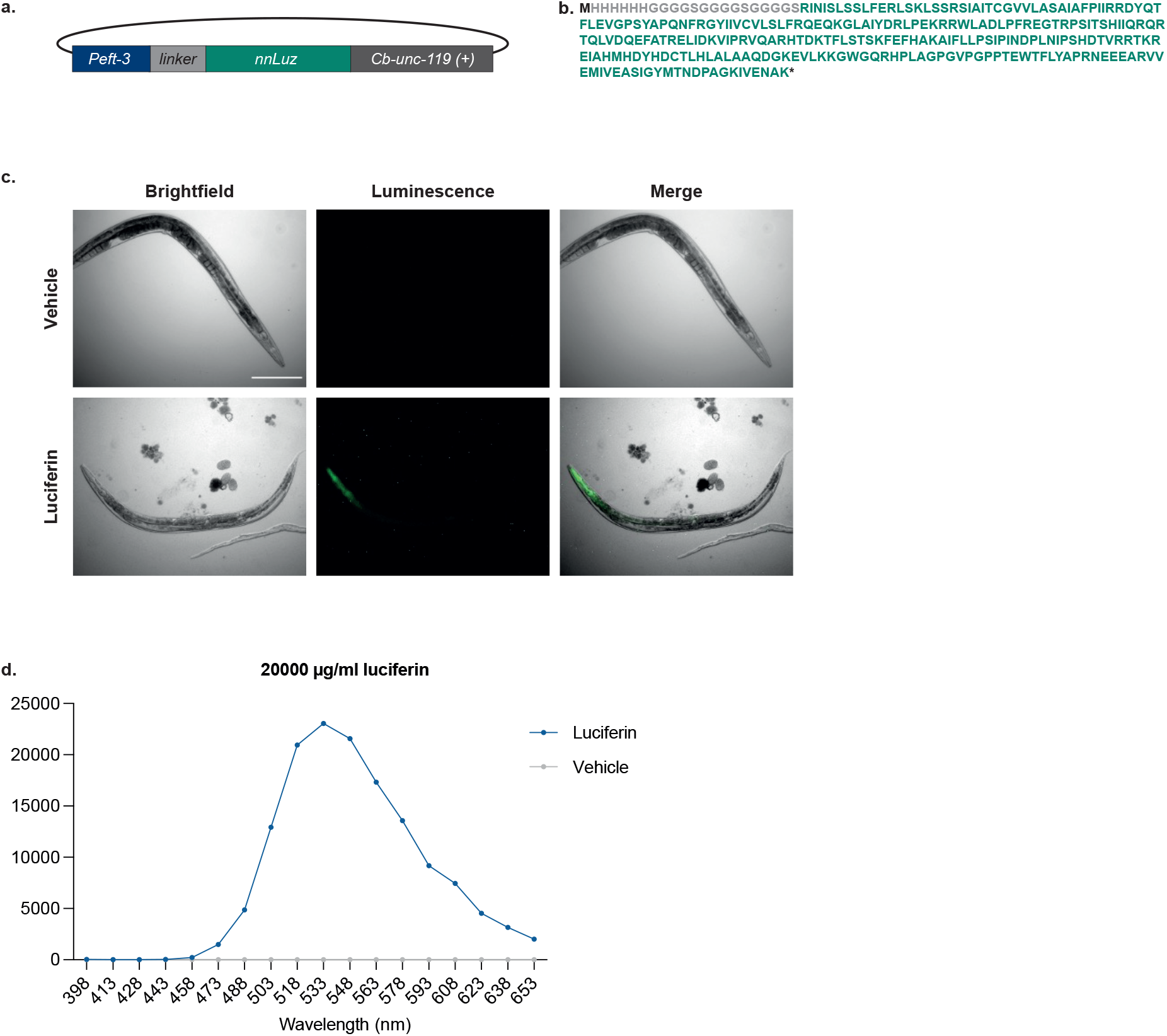
Luminescence emission from nnLuz expressing *C. elegans*. **a**. Schematic of the construct used to create a strain of *C. elegans* constitutively expressing nnLuz luciferase. Strain: ATGSi577, genotype: *fqSi11 [Peft-3 His Linker nnLuz; cb-unc-119(+)]II)*. **b**. Protein sequence of the linker and the Luciferase. **c**. Luciferase expressing worms were treated with 20000 µg/ml fungal luciferin or control (DMSO) and luminescence was observed in live worms (Scale bar, 200 µm). **d**. Emission spectra of luciferase expressing *C. elegans* treated with 20000 µg/ml fungal luciferin or control (DMSO).

## Methods

### Animal Maintenance

Mice were housed in a pathogen free facility in the following conditions: 21+/−2°C temperature; 45– 65% humidity; 12 h light-dark cycle with RM3 diet and water *ad libidum*. Tissues, wood blocks, and tunnels were used to enrich the environment. All animal procedures were performed under a UK Home Office Project Licence and Personal Licences in accordance with the British Home Office Animal (Scientific Procedures) Act 1986 and approved by the Imperial College AWERB committee.

### Cloning, CRISPR and Generation of *Cyp1a1-nnLuz* mESCs and mouse lines

The nnLuz_v3 gene sequence was provided by Light Bio, and codon-optimised for expression in mammalian cells. For overexpression experiments in HEK 293T cells, both *nnLuz_v3* and *iRFP670* DNA sequences were synthesized as dsDNA Gene Fragments (Twist Bioscience). DNA fragments were amplified by PCR and cloned into the pCAG vector (pCAG-GFP, Addgene plasmid 11150) through Gibson assembly method using the NEBuilder HiFi DNA Assembly Cloning Kit (E5520S, NEB) according to manufacturer instructions. Gene targeting of *nnLuz_v3-T2A-iRFP670* reporter into mouse *Cyp1a1* gene was performed using CRISPR in mESCs E14 with a sgRNA sequence (5’-TCTTCAGGCTTAGACTGTCC -3’) specific to cleave DNA 4 nucleotides downstream of the stop codon located in exon 7. The sgRNA sequence and its corresponding antisense complementary sequence were ordered as DNA oligos (Sigma) and cloned using BbsI restriction enzyme (Thermo) into the Cas9-GFP-containing plasmid pX458. pSpCas9(BB)-2A-GFP (pX458) was a gift from Feng Zhang (Addgene plasmid # 48138; http://n2t.net/addgene:48138; RRID: Addgene_48138)^23^. To induce reporter knock-in integration by DNA repair through homologous recombination, a donor DNA template containing 1 Kb of each homology arm was generated and used as a targeting vector. We also included a modified version of the internal ribosomal entry site sequence (IRES2) between the stop codon of *Cyp1a1* and the start codon of *nnLuz_v3* to allow independent protein synthesis of the reporter nnLuz_v3-T2A-iRFP670 polypeptide from the endogenous Cyp1a1 protein. In addition, a neomycin resistance cassette (*NeoR*) flanked by loxP sites was included in the donor DNA template for selection of mESC clones. Each component of the targeting vector were generated by PCR from existing DNA plasmids or, in the case of the homology arms, from genomic DNA from E14 mESC. After purification, PCR fragments were cloned into the targeting vector through Gibson assembly method using NEBuilder HiFi DNA Assembly Cloning Kit (E5520S, NEB) according to manufacturer instructions. CRISPR was performed on E14 mESC by co-transfecting both pX458 and targeting control plasmids using Lipofectamine 2000, followed by FACS enrichment of GFP positive cells 24 hours post transfection. GFP positive mESCs were seeded at very low density in 10 cm plates containing a layer of mitotically-inactivated mouse embryonic fibroblasts on 0.1% gelatin-coated dishes. After 1 week, mESC colonies were manually picked under the microscope and individually-expanded in the presence of Neomycin (150 μg/mL). Individual clones were screened by PCR and selected based on phenotypic similarities to parental non-transfected E14 mESC, such as growth rate and cell morphology. Karyotype was examined on selected clones prior to microinjection into blastocyst for mouse line generation.

Primers used for *Cyp1a1-nnLuz* mESCs and mice genotyping are outlined in table 1.

**Table 1:**
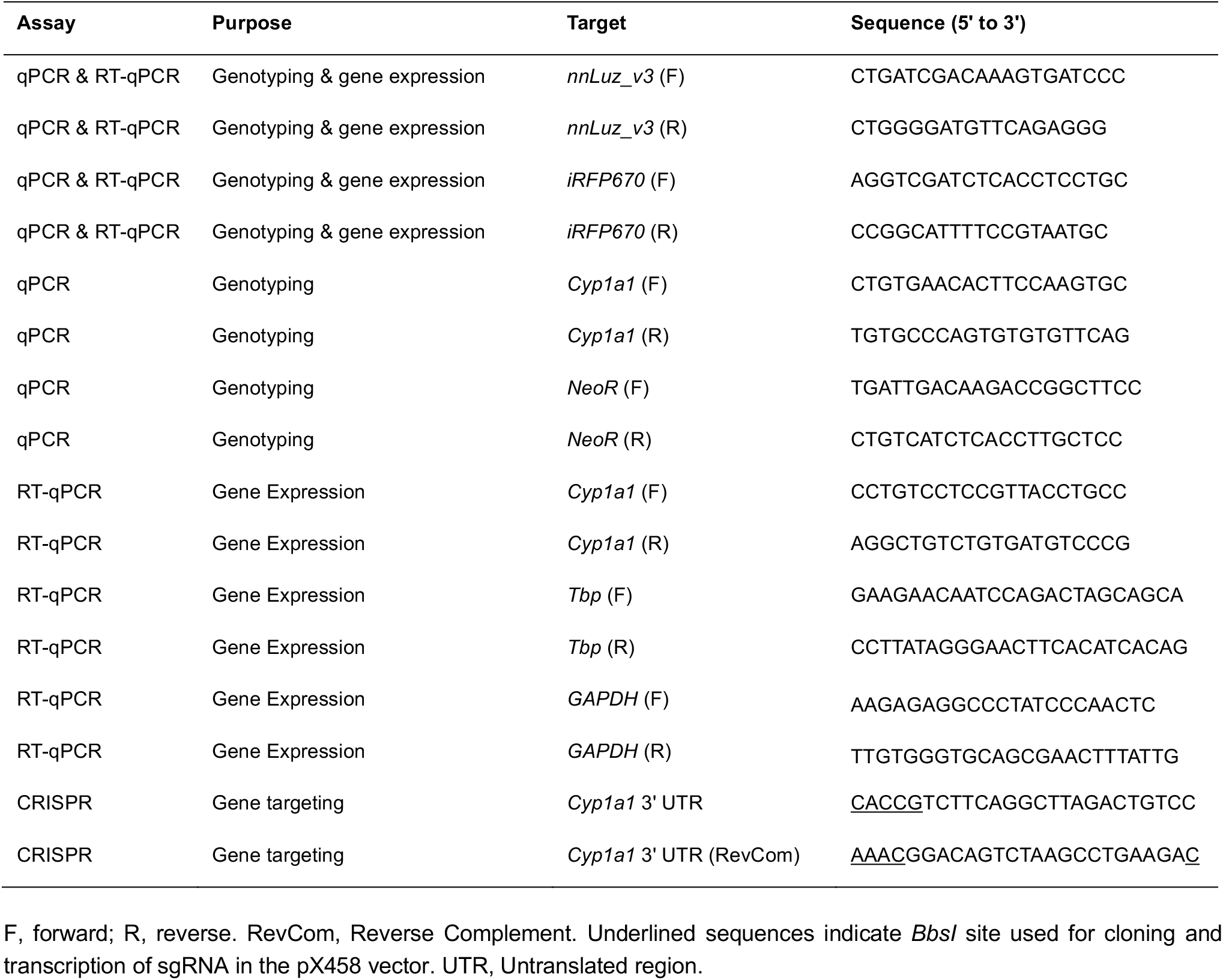
Primer and oligo sequences.

### Other animal lines

*Cyp1a1-Fluc* mice were generated by OzGene. A firefly luciferase (*Fluc*) gene was inserted into endogenous *Cyp1a1*, separated from the C-terminal region by a T2A sequence (see ^16^ for details).

### Mammalian cell culture and treatments

HEK 293T cells were grown in Dulbecco’s Modified Eagle’s Medium (Gibco), supplemented with 10% fetal bovine serum (Gibco), 1% penicillin-streptomycin (Gibco), 0.1 mM non-essential amino acids (Gibco) and 2 mM L-glutamine (Gibco). Cells were incubated at 37 °C with 5% CO_2_ and split every 3-4 days. ES-E14TG2a (E14) mESC line (129/Ola background) was used for gene targeting experiments to generate knock-in mESCs clones. mESC were cultured on a layer of mitotically-inactivated mouse embryonic fibroblasts on 0.1% gelatin-coated dishes with KnockOut Dulbecco’s Modified Eagle’s Medium (Gibco), supplemented with 15% fetal bovine serum (Gibco), 0.5% penicillin-streptomycin (Gibco), 0.1 mM non-essential amino acids (Gibco), 2 mM L-glutamine (Gibco), 0.1 mM 2-mercaptoetanol (Sigma) and 10^3^ U/mL of leukemia inhibitory factor (ESGRO, Millipore). Cells were incubated at 37 °C with 5% CO_2_ and split every 2–3 days. Plasmid DNA transfections on both HEK 293T cells and mESC were performed using Lipofectamine 2000 (Invitrogen) and Opti-MEM (Gibco), with cells in suspension in medium without penicillin-streptomycin, following manufacturer instructions. mESC were treated with 10 nM FICZ (BML-GR206-0100, Enzo) or DMSO (Sigma) as vehicle control.

### Fluorescence imaging and quantification

Fluorescent imaging of HEK 293T expressing pCAG-nnLuz-T2A-iRFP670 was performed on live cells cultured in 6-well plates and acquired using the EVOS M5000 Imaging system (Thermo), using the LED Cy5 (628/692 nm) light cube with 20X magnification.

### *in vivo* experiments

For all *in vivo* experiments involving *Cyp1a1-Fluc* or *Cyp1a1-nnLuz* animals, adult mice were first weighed and intraperitoneal (IP) injected with 3mC (213942, Sigma) dissolved in corn oil (Sigma) at 26.5 mg/kg. Corn oil was used as a vehicle control.

### Bioluminescence imaging

All bioluminescence imaging was performed using the IVIS Spectrum (Perkin Elmer) and Living Image software (version 4.3.1), using a stage temperature of 37°C.

Fungal luciferin was synthetised as described in ^24^. For imaging experiments luciferin was dissolved in DMSO to a concentration of 200 mg/ml, then diluted in dH_2_O to working concentrations on the day of use. D-luciferin (Perkin Elmer) was dissolved and stored in dH_2_O.

For bioluminescence imaging of HEK 293T cells and mESCs, fungal luciferin was diluted in DMEM or mESC medium, to achieve a final concentration of 650 µg/mL. Cells were imaged with the IVIS two minutes after exposure to fungal luciferin using field of view C, binning 4 and depth 0.5 for a total of 5 minutes.

For *in vivo* bioluminescent imaging experiments *Cyp1a1-Fluc* or *Cyp1a1-nnLuz* adult mice were IP injected with 3mC 5 h (IP) or ∼14 h (oral gavage) before luciferin exposure. Mice were IP injected with 0.15 mg/g D-luciferin (*Cyp1a1-Fluc*) or 0.15 mg/g fungal luciferin (*Cyp1a1-nnLuz*). For oral gavage, *Cyp1a1-nnLuz* adult mice were fed 0.15 mg/g fungal luciferin. Mice were anesthetized by isoflurane inhalation and imaged using the IVIS spectrum within 10-15 minutes of luciferin exposure. When assessing the kinetics of luminescence emitted from *Cyp1a1-Fluc* and *Cyp1a1-nnLuz* mice, consecutive images were taken across a 90–120-minute period and the animals remained anesthetized throughout. Adult mice were IVIS imaged using 30 second exposure, field of view C or D, binning 1 and depth 1.5. In the majority of cases the ‘open’ filter was used, capturing all emitted light. For generation of emission spectra, consecutive images were taken on the IVIS Spectrum using emission filters with wavelengths ranging from 500 to 700 nm with 20 nm band pass.

### RNA extraction and RT-qPCR

RNA was extracted using the RNeasy Mini Kit (Qiagen). For mESCs, cells were lysed using RLT buffer. Tissue samples were lysed in RLT buffer with a 5 mm stainless steel bead (Qiagen) using the TissueLyser II (Qiagen) for 4 min at 24,000 rpm. To digest fibrous tissue, heart samples were incubated with 10 μg/ml Proteinase K at 55°C for 1 h. Next, to remove debris, all tissue samples were centrifuged at top speed for 3 min and the supernatant retained. Total RNA was purified from the tissue supernatant or from cells lysed in RLT using the RNeasy Mini Kit according to the manufacturer’s instructions, including the on-column DNase digestion step using an On-column RNase-Free DNase Set (Qiagen).

RT-qPCR was performed using the QuantiTect SYBR Green PCR mix (Qiagen) in a CFX thermocycler (Bio-Rad) with 0.4 μM primers (sequences in Table 1). The amplification protocol consisted of 40 cycles of 94°C for 15 s and 60°C for 30 s. For each set of primers efficiency was calculated using the formula E = 10^−1/Slope^. Relative RNA levels were then determined using the formula EI^control Ct - sample Ct^/ EH^control Ct - sample Ct^, where EI = efficiency of primers of interest and EH = efficiency of the housekeeping primer set, *Tbp*.

### Luminescence Imaging of *C. elegans*

*Caenorhabditis elegans* expressing the *nnLuz* luciferase reporter (strain ATGSi577; genotype: fqSi11 [Peft-3 His Linker nnLuz; cb-unc-119(+)]II) were maintained on OP50-seeded NGM agar plates under standard conditions. Worms were picked from unsynchronized populations for luminescence measurements.

For luminescence assays, ten adult worms were transferred to individual wells of a black 96-well plate (Greiner 655096) containing M9 buffer (pH 9, adjusted with NaOH) supplemented with 20,000 µg/mL luciferin or DMSO control. Plates were incubated at 20°C for 30 minutes before luminescence emission spectra were recorded using a Tecan Spark plate reader. Following luminescence measurements, worms were transferred onto microscope slides with agar pads for imaging. Immobilization was achieved by adding a 2 µL droplet of 25 mM sodium azide. Imaging was performed using a Leica DMRB microscope equipped with a 10×/0.30 PH1 HC PL FLUOTAR objective lens and a Hamamatsu ORCA-ER digital camera (C4742-95) with a 10 second exposure and 4×4 binning.

### Statistics and reproducibility

Microsoft Excel was used for calculations and GraphPad Prism (version 10) used to create graphs and for statistical analysis. Graphs show the mean of experimental replicates with error bars showing standard error (SEM). Multi-group comparisons were tested using ANOVAs with Dunnett’s, or Šidàk’s multiple comparison tests. Specific details are provided in the figure legends.

